# An annual cycle of gene regulation in the red-legged salamander mental gland: from hypertrophy to expression of rapidly evolving pheromones

**DOI:** 10.1101/261495

**Authors:** Damien B. Wilburn, Richard C. Feldhoff

**Author notes:** Email addresses: DBW RCF.

## Abstract

Cell differentiation is mediated by synchronized waves of coordinated expression for hundreds to thousands of genes, and must be an exquisitely regulated process to produce complex tissues and phenotypes. For many animal species, sexual selection has driven the development of elaborate male ornaments, requiring sex-specific differentiation pathways. One such male ornament is the pheromone-producing mental gland of the red-legged salamander (*Plethodon shermani*). Mental gland development follows an annual cycle of extreme hypertrophy, production of pheromones for the ~2 month mating season, and then complete resorption before repeating the process in the following year. At the peak of the mating season, the transcriptional and translational machinery of the mental gland are almost exclusively redirected to synthesis of many rapidly evolving pheromones. Of these pheromones, Plethodontid Modulating Factor (PMF) has experienced an unusual history of disjunctive evolution: following gene duplication, positive sexual selection has diversified the protein coding region while the untranslated regions have been conserved by purifying selection. However, the molecular underpinnings that bridge the processes of gland hypertrophy, pheromone synthesis, and disjunctive evolution remain to be determined and are the focus of the present investigation. Using Illumina sequencing, we prepared a *de novo* transcriptome of the mental gland at six stages of development. Differential expression analysis and immunohistochemistry revealed that the mental gland initially adopts a highly proliferative, almost tumor-like phenotype, followed by a rapid increase in pheromone mRNA and protein levels. One likely player in this transition is Cold Inducible RNA Binding Protein (CIRBP), which selectively and cooperatively binds the highly conserved PMF 3′ UTR. CIRBP, along with other stress response proteins, have seemingly been co-opted to aid in mental gland development by helping to regulate pheromone synthesis. The *P. shermani* mental gland utilizes a complex system of transcriptional and post-transcriptional gene regulation to facilitate its hypertrophication and pheromone synthesis. The data support the evolutionary interplay of both coding and noncoding segments in rapid gene evolution, and necessitate study of the co-evolution of pheromone gene products along with their transcriptional and translational regulators. Additionally, the mental gland could be a powerful emerging model of regulated proliferation and subsequent resorption of a tissue, within the dermis, thus having potential links to skin cancer biology.

## Introduction

The processes of cellular differentiation and tissue remodeling are ubiquitous for multicellular organisms. The transition of cells from totipotency to terminal differentiation are the result of highly coordinated gene networks and changes in expression patterns [1]. Cell differentiation can be induced by a range of signals, including cytokines, hormones, cell contact, external stressors, and extracellular matrix composition [2-6]. Generally, gene expression is regulated through activation of specific transcription factors and/or post-transcriptional mechanisms (e.g. microRNAs and RNA-binding proteins) which permit the regulated expression of particular proteins [7, 8]. An extreme instance of targeted gene expression and differentiation is in the male ornaments of many animal species, such as ornate plumage in birds, vibrant coloration of fishes, and large antlers in deer [9, 10]. These elaborate ornaments are thought to arise from intense sexual selection imposed by female preferences and ultimately mate choice [11]. Such sexual selection can also influence molecular traits and has resulted in many reproductive proteins that evolve at extraordinary rates [12, 13]. For example, in the marine gastropod abalone, males produce an extraordinary number of sperm that overexpress the rapidly evolving protein lysin (up to ~1 g lysin per male abalone) that facilitates species-specific fertilization [14-16]; in fruit flies, males accessory glands synthesize complex mixtures of rapidly evolving seminal fluid proteins that restrict female re-mating by reducing her viability and survival [17, 18]; similarly, in *Pieris* butterflies, males produce enormous spermatophores (~13% of their body mass) that are encased in a nearly indestructible protein shell that slows spermatophore clearance and prevents female re-mating [19]. In these examples, however, the molecular mechanisms underlying the regulated expression of these unusual reproductive proteins are not fully understood.

Protein sex pheromones are another example of rapidly evolving reproductive proteins likely shaped by sexual selection through interaction with receptors in the other sex [13, 20]. For more than 60 million years, male plethodontid salamanders have utilized a system of nonvolatile protein courtship pheromones to regulate female reproductive behavior [21]. In the red-legged salamander (*Plethodon shermani*), during a courtship behavior known as tail-straddling walk, male salamanders privately deliver pheromones by “slapping” a large pad-like gland on his chin (the mental gland) to the female’s nares [22]. Pheromones diffuse into the female nasal cavity, bind to receptors on neurons in the vomeronasal organ, activate regions of the brain involved in pheromone response, and regulate female mating behavior [23-27]. Chemical analysis of the pheromone extract revealed two major components: Plethodontid Receptivity Factor (PRF) and Plethodontid Modulating Factor (PMF). PRF is a 22-kDa protein with sequence similarity to IL-6 cytokines [25], while PMF is a 7-kDa protein related to the highly diverse three-finger protein (TFP) superfamily that includes snake venom neuro- and cytotoxins, the complement receptor CD59, the human Ly6 antigen, the urokinase receptor uPAR, and the amphibian regeneration factor Prod1 [28]. When experimentally applied to female salamanders, both PRF and PMF altered the length of courtship time [25, 29-31]. Analysis by high performance liquid chromatography (HPLC) and mass spectrometry (MS) revealed multiple isoforms of both PRF and PMF; however, compared to 3 highly conserved PRF isoforms (>95% identity), individual male *P. shermani* expressed more than 30 diverse PMF isoforms (~30% identity) [32]. The ratios of different PRF and PMF isoforms are quite variable between male salamanders [33], and the source of isoform sequence diversity is primarily from gene duplication [32]. Examination of PRF and PMF sequences from 28 plethodontid species revealed that both genes have repeatedly experienced positive selection [34, 35]. Sampling from these many species by RT-PCR was facilitated by the unique quality that both PRF and PMF have unusually conserved, AU-rich untranslated regions (UTRs). The contrast is most striking for PMF: compared to the ~30% amino acid identity between isoforms, the average conservation for both the 5’ and 3’ UTRs is ~98%. We hypothesized that PMF genes have been subjected to disjunctive evolution: the coding regions of the many PMF gene copies have been under positive selection in order to expand the functional breadth of PMF as a pheromone, while purifying selection on the UTRs permitted coordinated, synchronized expression of the many PMF isoforms [32]. The mechanism(s) by which these UTRs mediate such expression remains unknown, but we postulated that RNA binding proteins were likely involved.

As with many elaborate male ornaments, the mental gland of male *P. shermani* is seasonally regulated. During the non-breeding season, it is mostly absent from male salamanders; however, presumably in response to elevated plasma androgens [36, 37], the gland hypertrophies over ~2 months and develops into a large pad-like structure solely dedicated to the production of protein pheromones (Fig 1). Once the gland has fully developed, PRF and PMF represent ~85% of the secreted protein [38]. Similarly, cDNA sequence analysis revealed that ~70% of the total mRNA coded for pheromones [39]. Following the end of the courtship season, the gland resorbs and a new one forms each subsequent year. It is noteworthy that surgical removal of the mental gland is followed by rapid wound healing and prevents gland regrowth in subsequent years (Lynne D. Houck and R.C. Feldhoff, personal communication), suggesting the existence of androgen-sensitive precursor cells embedded in the dermis. Given that the transcriptional and translational machinery of fully developed mental glands are directed almost exclusively towards pheromone production, there likely exist some earlier developmental phase characterized by greater mitosis and/or general growth to form the glandular structure. We hypothesized that the unusually conserved pheromone UTRs may be critical for regulating the transition from gland hypertrophication to pheromone synthesis. In this study, we applied transcriptome sequencing to characterize the *P. shermani* mental gland developmental profile about 3-week intervals from an early precursor stage to the active pheromone producing gland. We identified an RNA binding protein that binds to the highly conserved pheromone UTRs and likely contributes to the synchronized transition from gland hypertrophication to the overexpression of rapidly evolving pheromones during the short annual breeding season.

**Fig 1.**
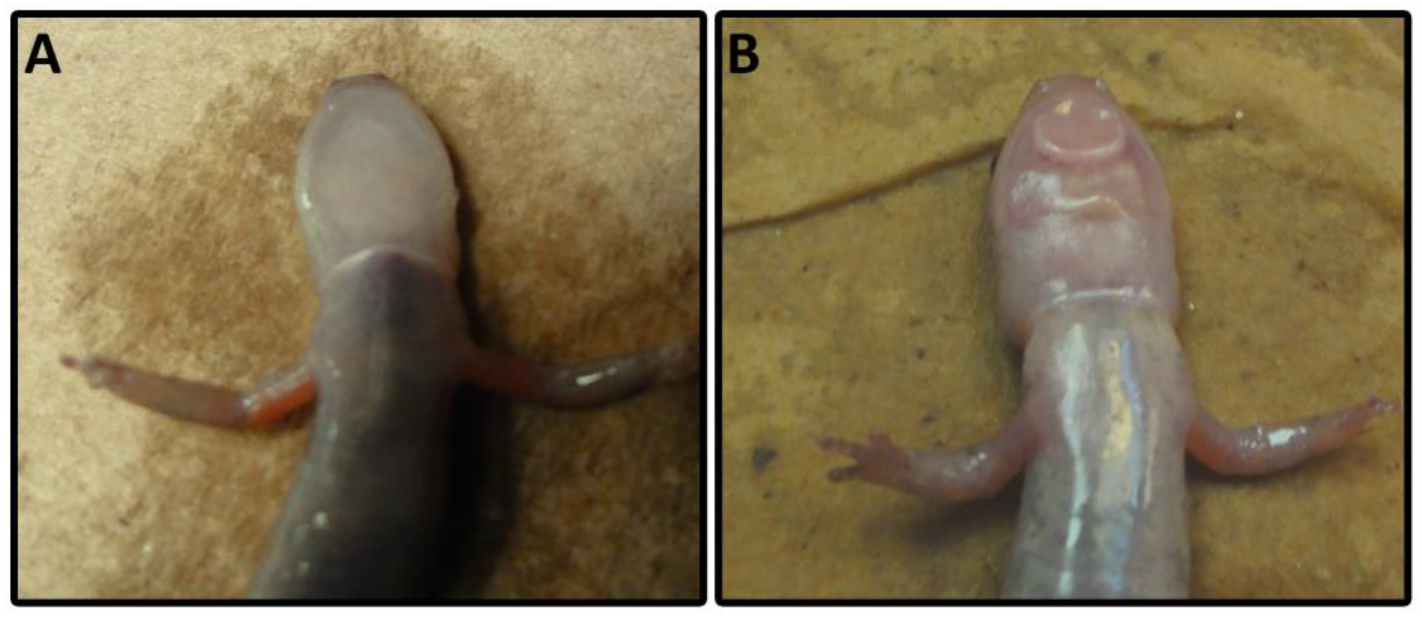
Mental gland hypertrophication. Comparison of male *P. shermani* from (A) late May (non-breeding condition) and (B) mid-August (breeding condition), with the mental gland being the large pad-like structure on the male’s lower jaw in panel B.

## Results

### Qualitative observation of mental gland development

In a previous study by Woodley [36], mental gland development was examined from mid-June to late August: during this period, plasma testosterone increased > 3 fold and mental gland volume significantly increased. Based on this work, we collected male *P. shermani* approximately every 3 weeks from late May through mid-September to span all phases of gland development. Our qualitative observations were similar to those of Woodley [36]. In late May, male salamanders had visibly different skin pigmentation near the mentum compared to females, including 2 of 15 males with extremely faint “outlines” of a mental gland. By mid June, 8 of 10 collected males had visible outlines and/or thin mental glands; and in mid July, all males collected had visible and protruding mental glands. At this mid-July point, pheromone was collected from 8 mental glands by incubation in an Amphibian Ringers buffer containing acetylcholine. Subsequent analysis by reverse-phase high performance liquid chromatography (RP-HPLC) revealed normal proportions of PRF and PMF, but with protein concentrations at ~33-50% of levels normally observed in mid-August. Glands from the last three time points (early August, late August, and mid September) were visibly well-developed and had normal levels of pheromone. In a second time series we collected animals from mid-June and early August, and observed similar progression of gland development, suggesting that mental gland development is tightly seasonally regulated in our study population. Histological analysis by hematoxylin/eosin staining showed that the glandular tissue was immediately under the epidermis (~2-3 cells thick). The mental gland was structured as cylindrically shaped bundles with nuclei localizing exclusively near the periphery. Strong eosin staining was observed in the center of these bundles during early August, contrasting with mid June where only nuclear staining was visible for the smaller, more tightly packed bundles (Fig 2A). Higher resolution images were obtained using confocal fluorescence microscopy: for early August, actin staining was strongest along the periphery (near the nuclei), with light, diffuse staining and a few small fibers visible in the eosin-stained space, whereas actin was only found adjacent to DAPI-stained nuclei in mid June (Fig 2B-C). Lectin staining suggested condensation and/or degradation of much of the ECM as the gland expanded (Fig 2B). Immunohistochemical labeling with anti-PRF produced a strong, punctate pattern throughout the eosin-positive space, reflective of secretory vesicles observed in scanning electron microscopy [40]. There was minimal PRF staining for mid June, both in intensity and volume (Fig 3). These data suggest that the mental gland initially forms as a tightly packed mass of cells with little cytoplasm, and upon induction of pheromone synthesis, cells swell with large volumes of pheromone, adopt a columnar shape, and the ECM condenses and/or degrades to support the enlarged cells.

**Fig 2.**
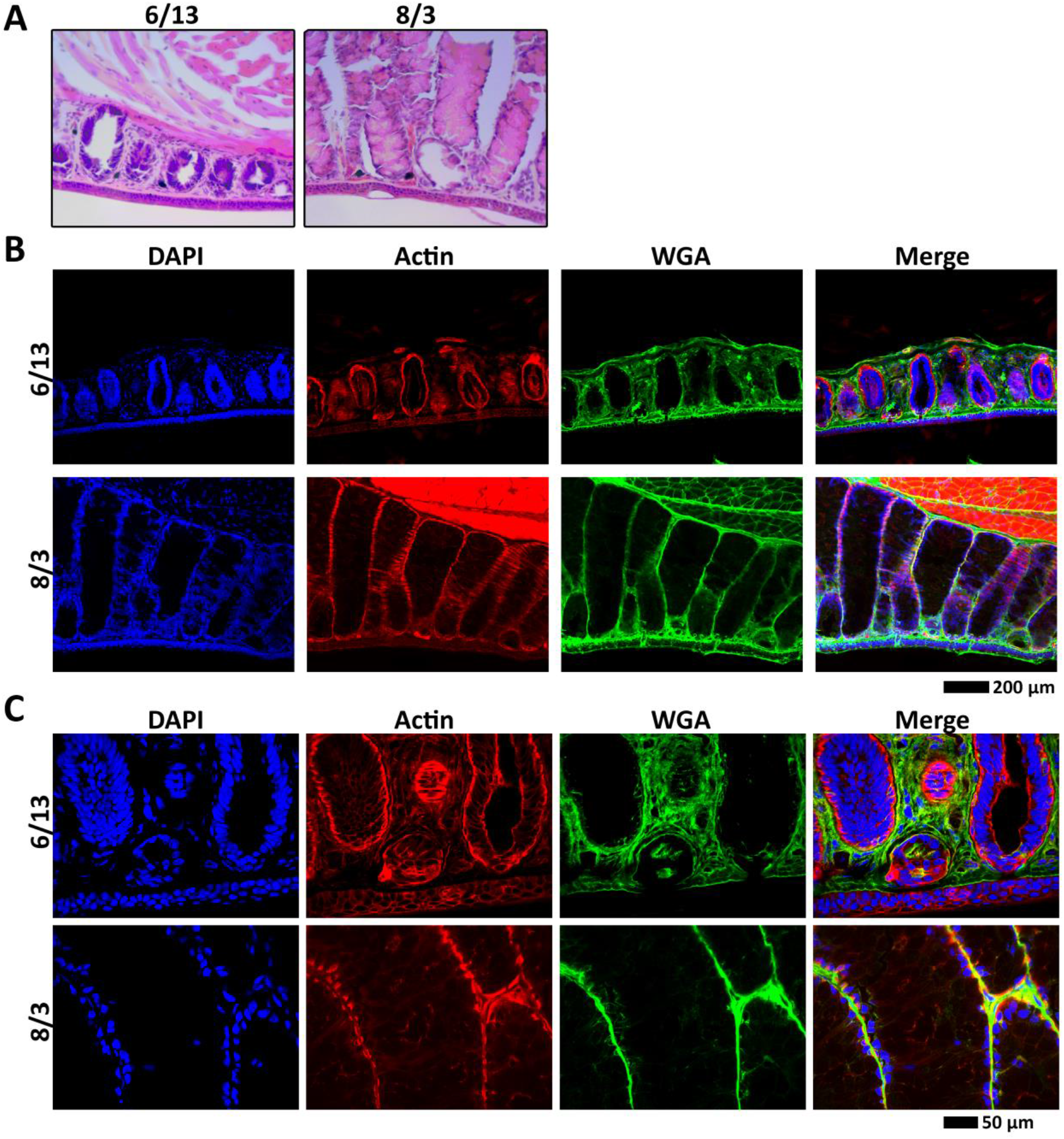
Mental gland histology. Comparison of mental glands from male *P. shermani* from two stages in mental gland development (mid June and early August) using (A) hematoxylin and eosin staining, (B-C) fluorescent confocal microscopy with dyes labelling the nucleus (blue), actin cytoskeleton (red), and ECM (green) at 10X (B) and 40X (C) magnifications.

**Fig 3.**
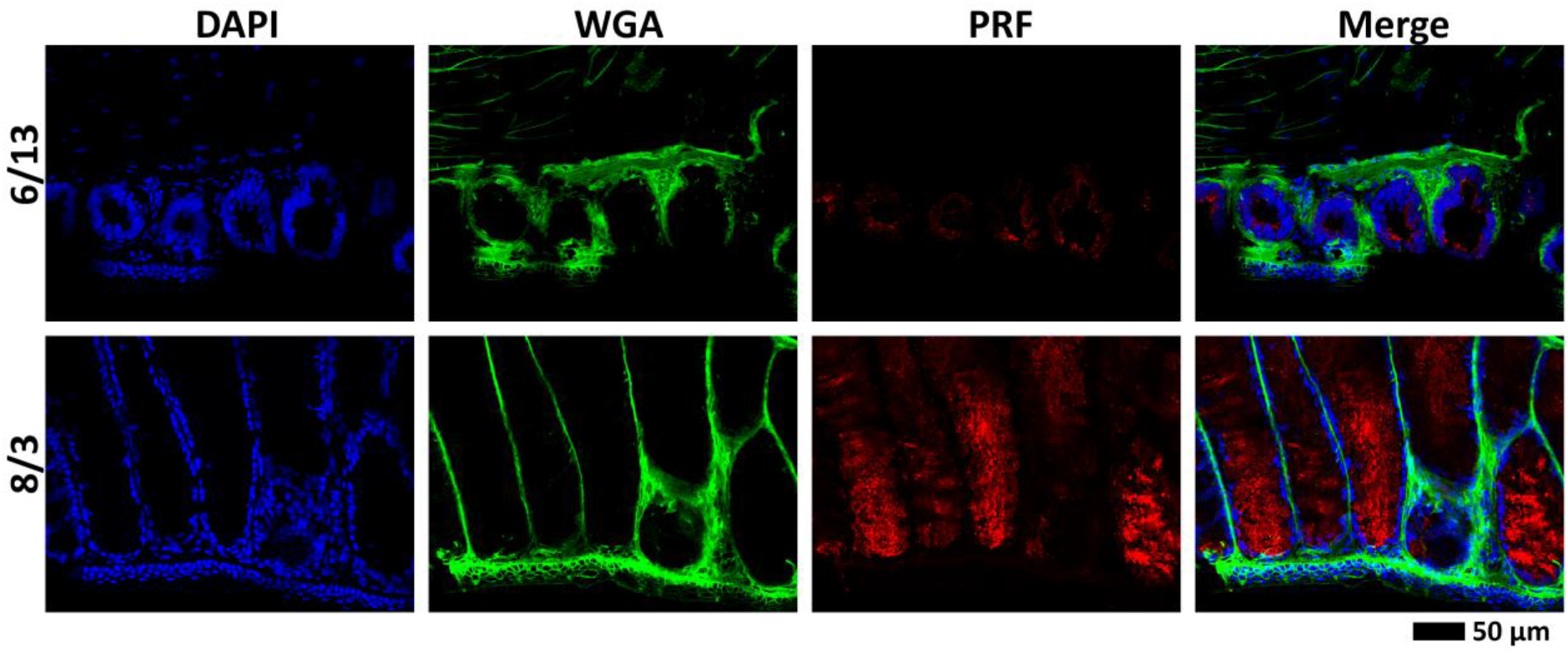
Pheromone immunohistochemistry. Comparison of pheromone expression and localization for mental glands at two stages of development (mid June and early August) by immunohistochemistry (using anti-PRF; red), with fluorescent dyes labelling the nucleus (blue) and ECM (green).

### Mental gland transcriptome and changes in gene expression

A transcriptome spanning mental gland development was constructed from the six time points spanning late May through mid-September. It should be noted that “mental gland” is used here to describe the tissue where the mental gland is normally found; for the earlier time points, this is largely comprised of skin and connective tissue. Interestingly, bands that correspond to PMF and PRF were visible in an agarose gel of full-length cDNA for all six time points, even in the early points when there was little-to-no pheromone protein (Fig S1). The intense ~850 bp band in the late May and mid June samples was later identified to encode a 15.3 kDa secreted protein with no significant BLAST hits in GenBank. A *de novo* transcriptome from Illumina reads was constructed using Trinity (Table S1), resulting in ~55,000 groups of reconstructed transcripts – herein referred to as genes for simplicity – which were annotated using several publicly available bioinformatics tools (results included in Supplementary Data 1). Nearly all assembled genes were detectable at all six points (at varying levels), and cluster analysis performed to identify genes with correlated expression patterns. Using a hierarchical approach, five major groups of genes were identified with different expression profiles and numbered 1 to 5 in order of decreasing gene count (Fig 4). Cluster 1 (which contained 87.7% of genes) showed maximum expression in May/June, and included many housekeeping genes (e.g. β-actin, GAPDH, PCNA, ferritin, ribosomal proteins). In relative terms, cluster 2 had the most stable expression patterns (~1-4X fold difference between time points), and included a range of genes from different biological pathways, including ribosomal proteins, lysosomal proteases, signal peptidase complex members, and lipid biogenesis enzymes. Cluster 3 included genes almost exclusively found in the earliest time point (with slightly elevated expression in the late August and mid September time points, possibly suggesting a cyclical response as the gland begins to resorb). Some of the most highly expressed genes in cluster 3 included ribosomal proteins (S6, S15, S17, S23, L14, L24, L32, L37a) and histone proteins (H1E, H2A, H3). Clusters 4 and 5 together include the genes most highly expressed in the later phases of gland maturation (0.8% of all genes). As expected, the majority of transcripts coded for pheromone, including PRF and PMF, but also (in lower abundance) many putative pheromones that were identified in other plethodontid species (natriuretic peptide, vasoactive intestinal peptide, sodefrin precursor-like factor, cysteine rich secretory protein) [39, 41, 42]. Included in these sequences was a predicted protein related to the tissue inhibitor of metalloproteinase (TIMP) family that included an extraordinarily long 3’ UTR (~3700 nt); by mass spectrometry, this sequence matched a protein previously termed C3 that constitutes ~10% of the pheromone extract [33]. The function of this protein is still unknown, but given this new information as to its likely homology, we now refer to it as Plethodontid TIMP-like Protein (PTP) [20, 41]. Multiple other protease inhibitor-like proteins were identified, included cystatin C and multiple Kazal-type inhibitors. Clusters 4 also included several retrotransposon and reverse transcriptase-like sequences. The biological importance of these sequences is unclear, but may explain the presence of intron-less processed PMF pseudogenes in the *P. shermani* genome [32]. Three other proteins of interest in cluster 4 included acetylcholinesterase (AChE), nuclear protein 1 (NP1), and vascular endothelial growth factor (VEGF). Incubation with acetylcholine is the standard methodology by which pheromones are extracted from the mental gland [25, 43]; under normal biological conditions, co-secretion of AChE would allow for tightly controlled pheromone release [44]. VEGF, an angiogenic factor, may be necessary for mental gland maintenance: as the mental gland enlarges, the cells closest to the dorsal surface will be distant from dermal capillaries and may not receive sufficient nutrition without recruitment of new blood vessels. Related, NP1 classically functions in chromatin remodeling as part of stress responses (such as nutrient starvation) to prevent apoptosis [45,46]. In mental gland development, NP1 is one of the few genes to steadily increase in mRNA expression over the time course, and may play an important role in ensuring that the gland persists throughout the courtship season after it has transitioned to pheromone synthesis. As a quality control, qRT-PCR was performed for 16 genes of interest (Fig S2), and similar expression patterns were observed between RNASeq- and qPCR-based estimates, with no significant biases from gene or time point.

**Fig 4.**
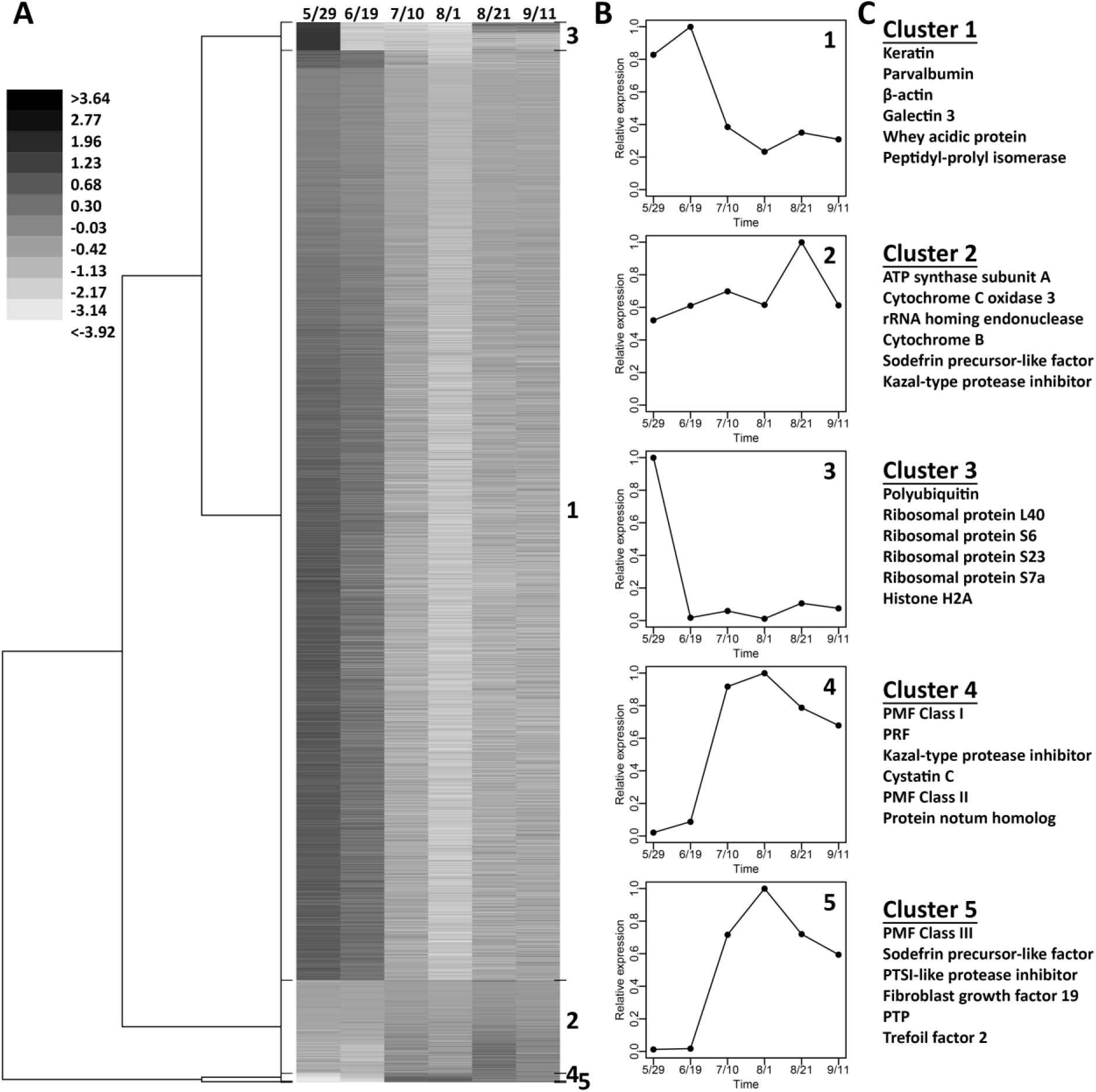
Cluster analysis of mental gland gene expression. (A) Cladogram representing the ~55,000 “genes” organized into 6 clusters. Shades of grey represent log fold changes between time points. (B) Line graphs of cluster means vs time. (C) The six most abundant genes in each cluster (at the time point with the highest expression levels per cluster).

### Cold Inducible RNA Binding Protein (CIRBP) binds the PMF 3’ UTR

The unusual pattern of disjunctive evolution for PMF – rapidly evolving coding sequence with conserved UTRs – led us to hypothesize that the UTRs may recognize RNA binding proteins (RNA-BPs) to coordinate the expression of the many diverse PMF isoforms [32]. While no RNA-BPs were detected as differentially expressed with a 5% false discovery rate, we found one highly abundant RNA-BP by manual inspection: Cold Inducible RNA-BP (CIRBP), which represented ~0.13% of transcripts in mid June. Analysis by qRT-PCR confirmed differential mRNA expression over the six time points, with maximum expression at mid June (Fig S2). To test for biological activity *in vitro*, recombinant CIRBP fused to the enhanced cyan fluorescent protein (rCIRBP/ECFP) was expressed in *E. coli*, and electrophoretic mobility shift assay (EMSA) experiments were performed with rCIRBP/ECFP and *in vitro* transcribed RNA. rCIRBP/ECFP was titrated against a nearly full length PMF 3’ UTR (nucleotides 26-667), and a very clear shift was observed in both the RNA and protein bands (Fig 5). Interestingly, there was visible RNA smearing at lower concentrations of rCIRBP/ECFP, suggesting possible dissociation of the RNA-protein complex during electrophoresis. Simultaneously, the altered position of the RNA band in the presence of greater rCIRBP/ECFP suggested a non-equimolar stoichiometry. To narrow the potential binding sequences for CIRBP, four overlapping ~250 nt segments of the PMF 3’ UTR were prepared, along with a similar sized segment of the keratin 3’ UTR as a negative control. When these five different RNAs were analyzed by EMSA, all showed visible gel shifts in the presence of increasing rCIRBP/ECFP, yet the overlap between RNA/protein was most intense in the PMF 3’ UTR 26-288 and 99-368 fragments (Fig S3). Variability in band number and intensity in zero-protein control lanes suggested different degrees of RNA secondary structure between the different sequences, which may have had an impact on CIRBP binding. Specificity of CIRBP towards the PMF 3’ UTR was assessed using a competition assay where fluorescently tagged versions of the different RNA molecules were observed in the presence and absence of excess unlabeled RNAs. With PMF 3’ UTR 99-368, addition of a 100-fold excess of unlabeled PMF 3’UTR 99-368 eliminated the gel shift, while a 100-fold excess of unlabeled keratin 3’ UTR only reduced the gel shift to a smear (Fig 6A). These data suggest that CIRBP has relatively greater affinity for the PMF 3’ UTR, with some lower affinity for other RNA molecules. In a similar competition assay using fluorescent PMF 3’ UTR 26-667, 100X unlabeled RNA was added for three different lengths of the PMF 3’ UTR (26-288, 26-565, 26-667) and keratin 3’ UTR. Only the full length PMF 3’ UTR 26-667 was able to fully eliminate the observed gel shift, suggesting a positive correlation between RNA length and strength of CIRBP binding (Fig 6B).

**Fig 5.**
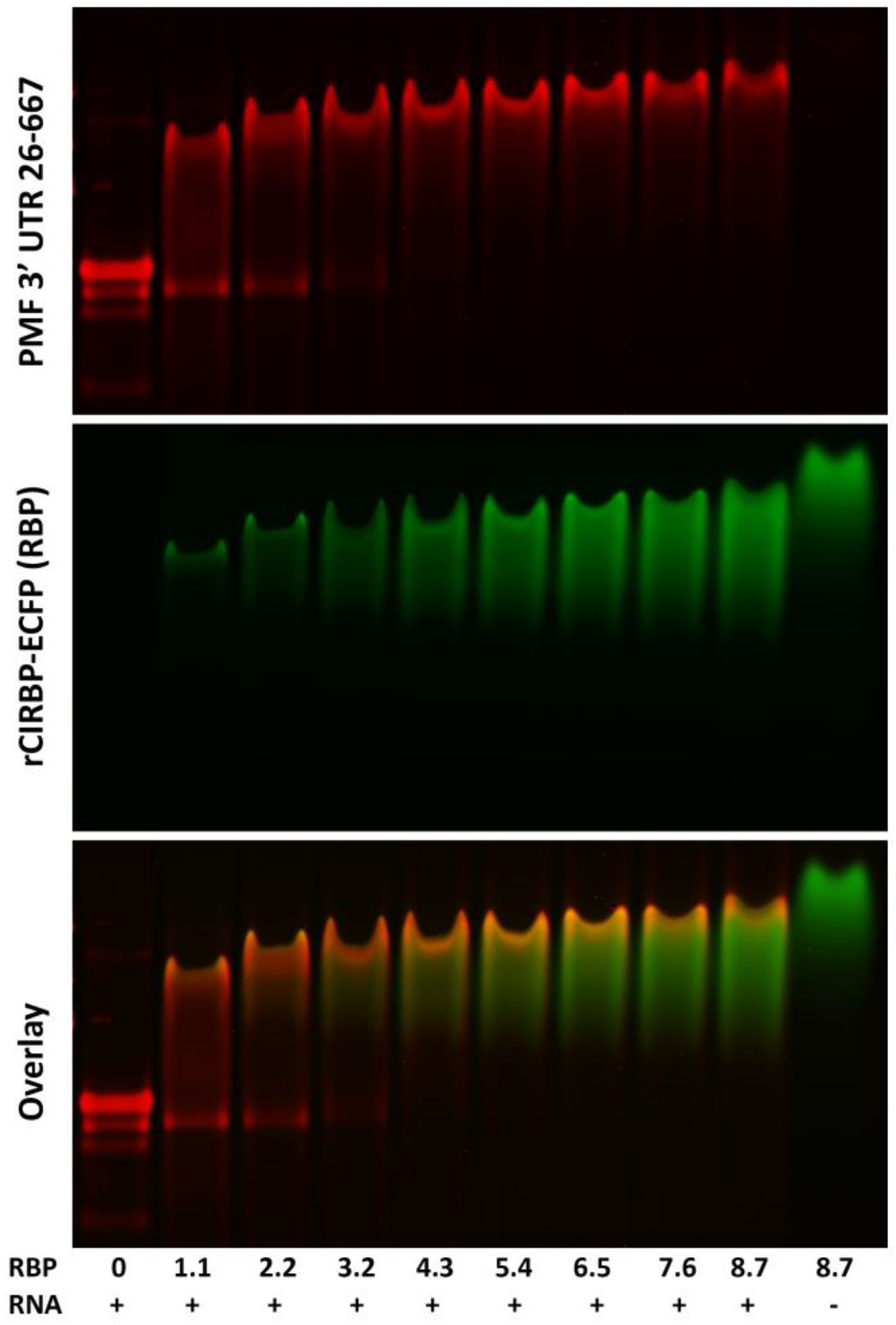
CIRBP – PMF 3’ UTR interaction. EMSA using a constant amount of PMF 3’ UTR *in vitro* RNA (200 ng) with increasing concentrations of rCIRBP/ECFP (μM; RBP), and a protein-only control. Protein fluorescence was detected by ECFP (green), and RNA was stained using Sybr Green II (red).

**Fig 6.**
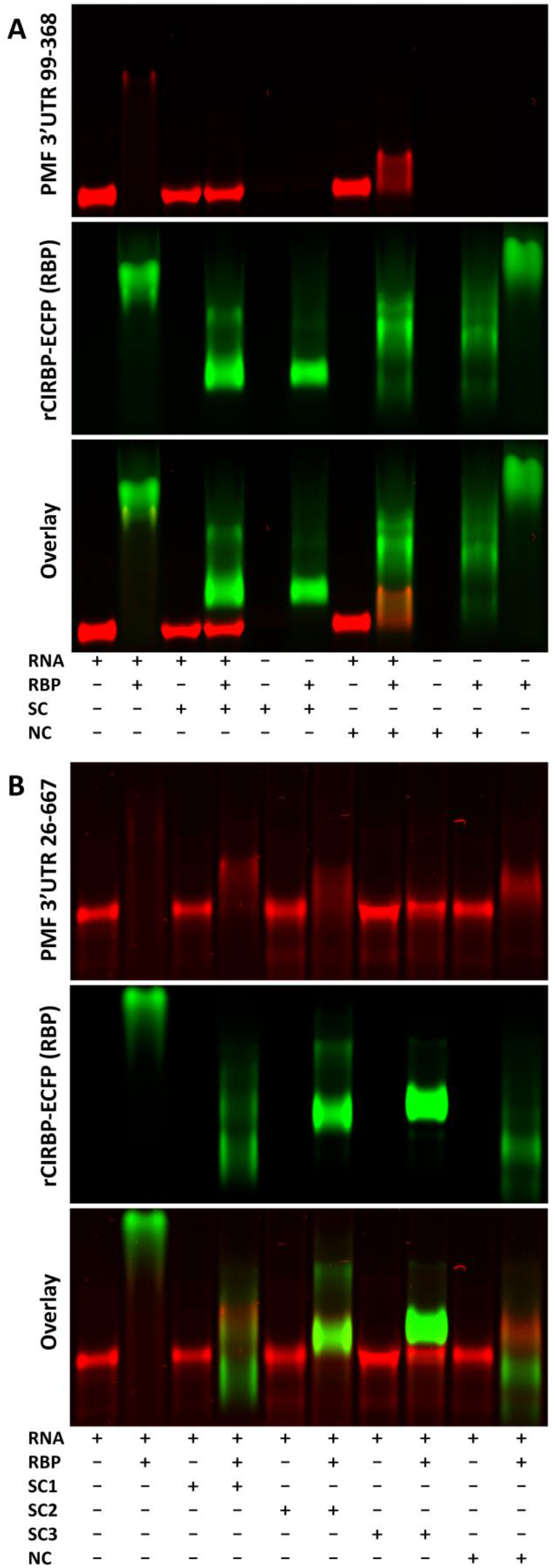
Competition EMSA with CIRBP. (A) EMSA between TAMRA-labeled PMF 3’ UTR 99-368 RNA (30 ng; red) and rCIRBP/ECFP (1.5 μg; RBP; green), with competition using 100X (3 μg) of a specific competitor (SC; unlabeled PMF 3’ UTR 99-368) or a non-specific competitor (NC; unlabeled Keratin 3’ UTR 205-441). (B) EMSA with TAMRA-labeled PMF 3’ UTR 26-668 RNA (30 ng; red) and rCIRBP/ECFP (1.5 μg; RBP; green), with competition using 100X (3 μg) of four different unlabeled competitors: SC1 = PMF 3’ UTR 26-288, SC2 = PMF 3’ UTR 26-565, SC3 = PMF 3’ 26-668, or NC = Keratin 3’ UTR 205-441.

### Dynamics of CIRBP-PMF 3’ UTR interactions

CIRBP contains two structural domains: a N-terminal RNA recognition motif (RRM) and a C-terminal glycine-rich, low complexity domain (LCD). Studies on the human homolog of CIRBP suggested that both domains bind RNA, with the RRM and LCD having specific and non-specific interactions, respectively [47]. For *P. shermani*, each domain was expressed as a separate fusion protein to enhanced cyan fluorescent protein (rCIRBP-RRM/ECFP and rCIRBP-LCD/ECFP), and neither domain in isolation induced strong gel shifts with fluorescent PMF 3’ UTR (99-368) (Fig S4). A small amount of smearing that occurred in the highest concentrations of rCIRBP-LCD/ECFP, suggested a weak interaction. As similar smearing was observed at lower concentrations with rCIRBP/ECFP, EMSAs with rCIRBP/ECFP and PMF 3’ UTR 99-368 were repeated with and without formaldehyde treatment to crosslink protein-RNA complexes. Crosslinking successfully reduced the amount of visible RNA smearing in the gel (Fig S5), suggesting that under sufficiently low stoichiometry, rCIRBP/ECFP (and likely rCIRBP-LCD/ECFP) forms unstable complexes that readily dissociate during electrophoresis.

When using fluorescently labeled RNAs, interaction with CIRBP caused fluorescence quenching of bound RNA – likely due to shielding of the fluorophores attached to the uracil bases (Fig 6). This fluorescence quenching was exploited to estimate binding affinities of all three CIRBP constructs to PMF 3’ UTR 99-368 in solution (Fig 7). While there was no detectable fluorescence quenching with rCIRBP-RRM/ECFP, both rCIRBP/ECFP and rCIRBP-LCD/ECFP yielded sigmoidal curves characteristic of cooperative binding. Fitting of this data to the Hill equation by nonlinear regression yielded both significant association constants (K_A_) and Hill coefficients (n), but with rCIRBP-LCD/ECFP having both lower affinity (higher K_A_) and weaker cooperativity (lower n) compared to rCIRBP/ECFP. Therefore, there likely exists synergism between the two domains to promote binding to the PMF 3’ UTRs.

**Fig 7.**
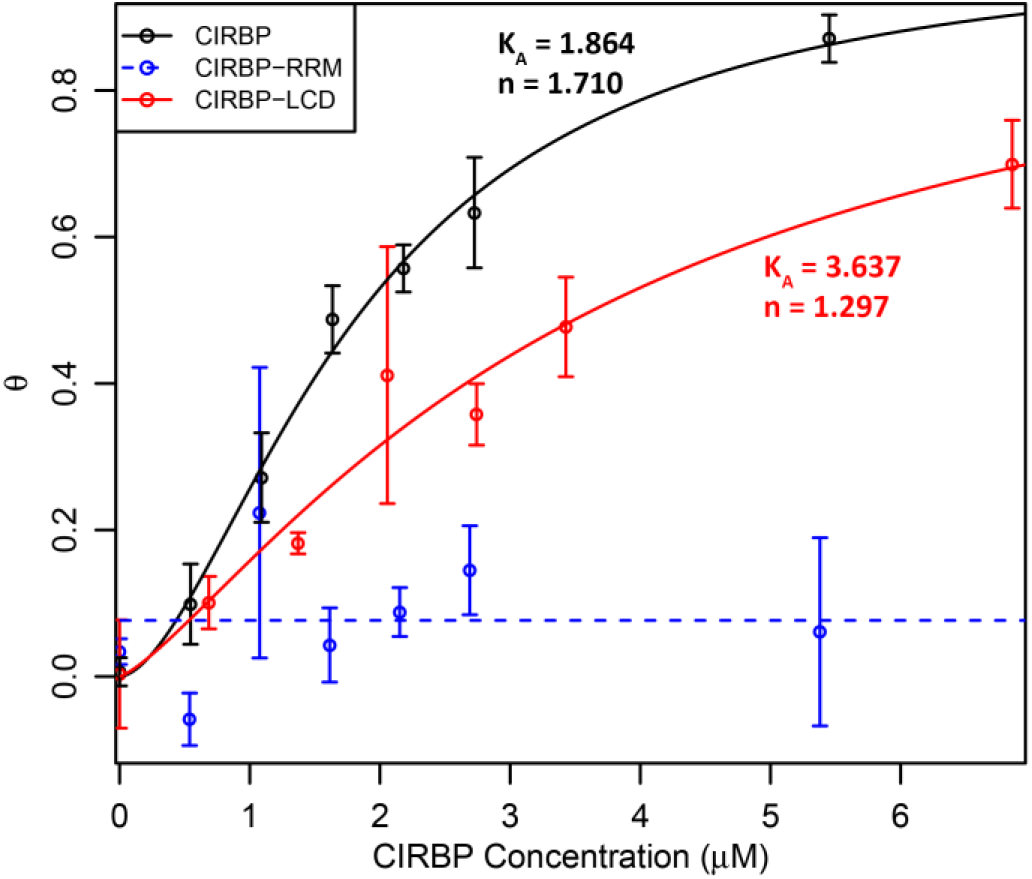
CIRBP titration curve. TAMRA-labeled PMF 3’ UTR 99-368 RNA was titrated with increasing concentrations of rCIRBP/ECFP (black), rCIRBP-RRM/ECFP (blue), and rCIRBP-LCD/ECFP (red), and binding measured by fluorescence quenching. Data were fit to the Hill equation by nonlinear modeling to obtain measures of binding affinity (K_A_) and cooperativity (n). No significant change was detected for rCIRBP-RRM/ECFP, denoted by a dashed line.

Recent models of other RNA-BPs with LCDs suggested that, upon binding to a proper catalyst, unstructured LCDs adopt regular β-sheet structure that drive aggregation and formation of stress granules, processing bodies, or other macromolecular RNA-protein complexes [48]. To test if the PMF 3’ UTR may be acting as such a catalyst, rCIRBP/ECFP was analyzed by circular dichroism (CD) and titrated with increasing amounts of PMF 3’ UTR 26-667 (Fig 8). There was a detectable increase in the CD absorbance, particularly near ~215 nm where β-sheet can be measured. While the percentage change is relatively small, the majority of CD signal likely originates from ECFP (a highly structured β-barrel that comprises ~60% of the fusion protein), and since CD only reports on the average secondary structure content, the “induced” β-sheet in CIRBP may only be occurring in a small proportion of the available molecules. These data support that binding of CIRBP to the PMF 3’ UTR promotes a conformational change and increased secondary structure, likely in the LCD shifting from random coil to ß-sheet.

**Fig 8.**
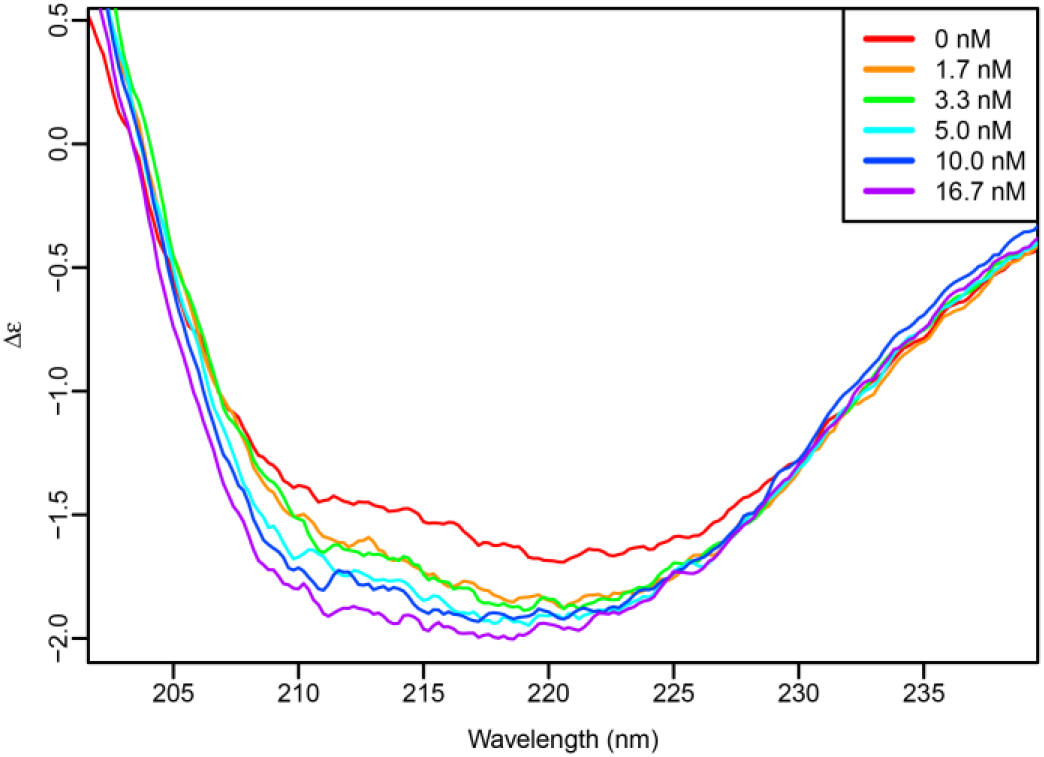
PMF 3’ UTR induces secondary structure changes in CIRBP. CD spectra of rCIRBP/ECFP with increasing concentrations of PMF 3’ UTR 26-668 RNA, with changes in CD suggesting higher levels of secondary structure (likely ß-sheet).

### CIRBP expression and correlation to PMF in vivo

Polyclonal antibodies to rCIRBP/ECFP were prepared and affinity purified specifically against CIRBP-RRM. Immunohistochemical staining using anti-CIRBP-RRM revealed that CIRBP protein was localized to the cytoplasm, and more abundant mid June compared to early August. The staining was uniformly distributed, uncharacteristic of cytologically visible RNA granules such as stress granules, Cajal bodies, or nuclear speckles (Fig 9). CIRBP protein levels for individual mental glands were estimated by western blot analysis. However, multiple bands were observed near and below the approximate 17 kDa expected molecular weight (Fig 10A-D). We postulated that these bands were CIRBP degradation products generated by a protease sequentially processing the C-terminal LCD. In support of this hypothesis, when proteins from immuno-pulldown were analyzed by mass spectrometry, peptides were identified that fully span the N-terminal RRM (Fig 10B). Addition of either a broad protease inhibitor cocktail or 0.1 mM iodoacetamide (to specifically inhibit cysteine proteases) limited the extent of degradation, but did not completely ablate it (Fig 10A). The same samples were examined by western blot analyses over multiple days, and even without a protease inhibitor, degradation was incomplete (data not shown). Thus, some of this visible degradation may be a result of natural CIRBP turnover and perhaps play a biological role. Interestingly, neither inhibitor altered the number of observed bands, but rather their relative intensities, such that a single protease may be contributing to both natural and experimentally-imposed degradation. While we were unable to determine the specific protease involved, multiple lines of evidence suggested that it may be cathepsin S: (1) iodoacetamide performed as well or better than the diverse mixture of protease inhibitors, implicating a cysteine protease; (2) most of the identified mental gland proteases in the transcriptome were lysosomal, with cathepsin S having the highest expression; (3) in contrast to most lysosomal proteases, cathepsin S is active only at near-neutral pH, and the lysis buffer was at pH 8; (4) the CIRBP LCD is enriched for the cathepsin S target sequences (aliphatic or aromatic residues followed by Gly [49], specifically YG in the CIRBP-LCD), and cleavage at these sites would generate proteins of similar molecular weight to those observed by western blot (Fig 10C).

**Fig 9.**
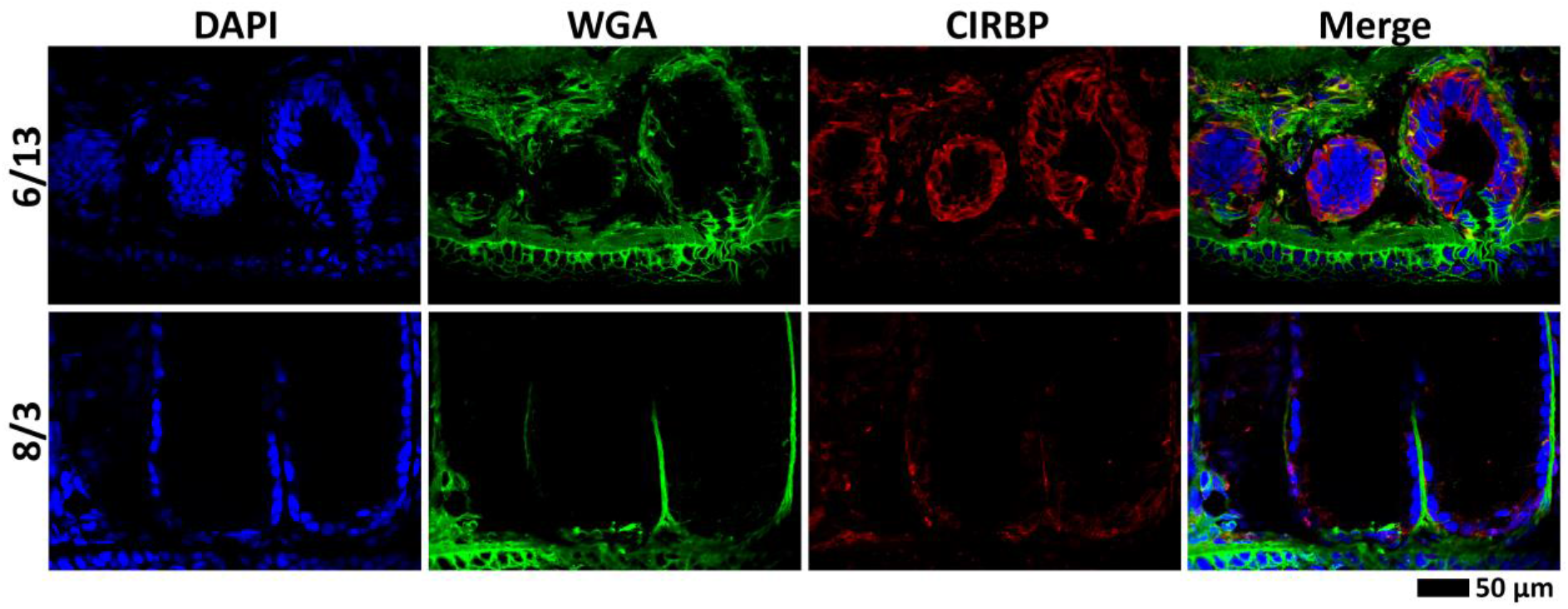
CIRBP immunohistochemistry. Comparison of CIRBP expression and localization for mental glands at two stages of development (mid June and early August) by immunohistochemistry (using anti-CIRBP-RRM; red), with fluorescent dyes labeling the nucleus (blue) and ECM (green).

**Fig 10.**
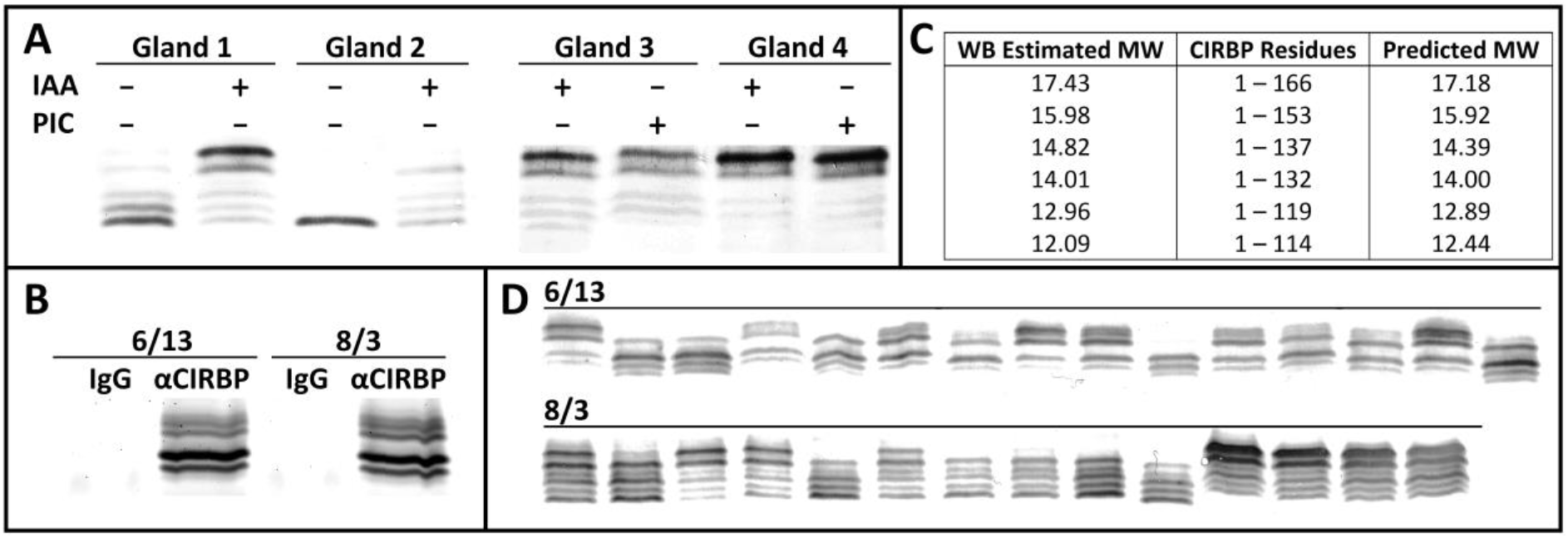
CIRBP protein analysis. (A) Western blot analysis comparing CIRBP degradation with and without different protease inhibitors. Mental glands were dissected into approximate halves, and incubated with RIPA extract containing no protease inhibitors, a commercially available protease inhibitor cocktail (PIC), or 100 μM iodoacetamide (IAA). (B) SDS-PAGE with SYPRO Ruby stain of immunopulldown products using either a rabbit IgG mixture vs anti-CIRBP-RRM for two time points in gland development. (C) Estimated molecular weights (in kDa) for different CIRBP degradation products compared with masses predicted by cleavage of C-terminal Cathepsin S sites (YG). (D) Western blot analysis demonstrating the diversity of CIRBP abundance and degradation state for individual mental glands from mid June (5 μg total protein) vs early August (30 μg total protein) when extracted in the presence of IAA.

To ascertain a potential role for CIRBP in regulating mental gland development and PMF synthesis, several variables were correlated from individual mental glands collected at two time points (mid June and early August). Pheromone was extracted by incubation in acetylcholine and protein concentrations measured. Because PMF consistently comprises ~50% of the total pheromone [33], we used pheromone concentration as a proxy for PMF protein expression (P_PMF_). Following pheromone extraction, mental glands were stored in RNAlater, and later dissected into two approximately equal halves, permitting independent isolation of total RNA and cellular protein. Using standardized amounts of cellular protein, CIRBP expression was measured by western blot and densitometry for both total CIRBP (*_PTotal-CIRBP_*) and intact CIRBP (P*_Intact-CIRBP_*) (Fig 10D). Using total RNA, expression levels were measured for three genes by qRT-PCR: PMF (R*_PMF_*), CIRBP (R*_CIRBP_*), and cathepsin S (R*_Cath_*). Using these 6 variables plus a time covariate (mid June vs early August), a series of nested MANOVAs were performed to identify potentially meaningful correlations (Table S3). Time was a significant covariate for all variables, and the only significant factor for all three measured mRNA levels. Total CIRBP protein (including degradation products) increased proportionally to CIRBP mRNA levels, but with a higher ratio in June compared to August. Total CIRBP was the best predictor for intact CIRBP levels, however, there were significant interaction terms between *time*/R*_Cath_* and *time*/R*_Cath_*/R*_CIRBP_*/P_Total-CIRBP_ such that only in June, intact CIRBP levels decreased as Cathepsin S mRNA levels increase (both independently and in proportion to total CIRBP levels). Finally, while neither CIRBP protein variable affected PMF mRNA levels, pheromone protein levels were negatively correlated with intact CIRBP (reflected by the *time*/P*_Intact-CIRBP_* interaction term, since no pheromone protein detected in June). While some of these correlations may be a consequence of natural gland progression in response to other unmeasured variables, these data provide additional evidence to support that Cathepsin S regulates steady-state levels of CIRBP. As intact CIRBP was negatively correlated with PMF protein levels but not mRNA levels, we hypothesize that CIRBP may be acting as a translational repressor.

## Discussion

The process of mental gland development is an exciting example of sex-specific differentiation and organogenesis that yields an elaborate male ornament which expresses numerous rapidly evolving reproductive proteins. During the non-breeding season, the mentum of male *P. shermani* appears as normal skin and is visually indistinguishable from that of a female. A particularly fascinating feature of this system is the annual nature of accelerated growth followed by natural resorption, with the first half of the process sharing several qualities with tumorigenesis. The presented molecular and histological data support our original hypothesis that mental glands begin in a highly mitogenic state, and then transition into a veritable pheromone factory. Cells first rapidly divide and proliferate, evidenced by elevated PCNA levels at the earliest time points in gland development (Fig S2). As these cells divide, many housekeeping genes (representing over 80% of the expressed genes) are activated and construct the basic architecture of the gland (Fig 4). The overall glandular structure seems to consist of many dozens to hundreds of cells surrounding an open lumen with a channel leading to the epidermis (Fig 2). At early stages of development, these cells consist volumetrically of little more than a nucleus, characteristic of rapid mitosis and further mimic a tumor-like structure [50-52]. While describing the early mental gland as “tumor-like” may be allegorical, there is a reasonable degree of analogy: the mental gland seemingly “invades” the dermis, compresses and/or degrades the ECM, rapidly divides to form a larger structure, and eventually releases angiogenic factors to increase blood supply to extend its viability. While ECM composition was only qualitatively examined through lectin staining (Figs 2, 3, 9), it is noteworthy that several different protease inhibitor families (TIMPs, cystatins, Kazal-type serine protease inhibitors) are overexpressed at later stages of development. Many of these proteins have signal peptides, and may be secreted to inhibit proteases that are initially required for mental gland expansion, but must be later inactivated to prevent cell degradation. Notably, Plethodontid TIMP-like Protein is part of the pheromone mixture when extracted with acetylcholine [33], and is presumably released when male salamanders slap to deliver pheromone. It is unclear if the other proteins are released in the same manner, or secreted in response to other stimuli. Alternatively, these protease inhibitors may be necessary to protect synthesized pheromones from premature degradation. What makes this tumor-like phenotype particularly interesting is that, unlike normal cancerous tissue, the mental gland completely resorbs into a non-precursor state following the courtship season. Detailed molecular characterization of both the proliferation and resorption processes could have powerful implications towards cancer biology and identifying potential factors with which to reprogram tumor cells.

For the gland to transition towards pheromone synthesis, extensive changes in gene expression are required, including overexpression of both PRF and PMF mRNA. While the proportions changed, nearly all genes in the transcriptome were detectable at all 6 time points. On average, housekeeping genes in Cluster 1 were still represented at ~33% of maximum levels after the upregulation of pheromone synthesis. Despite minimum pheromone translation at early time points (Fig 3 and Table S3), PRF and PMF together comprised ~1.5% and ~10% of all mRNA in May and June, respectively. One possibility may be that androgen stimulation uniformly activates transcription of all mental gland-associated genes. Subsequently, additional variables such as chromatin remodeling and mRNA stability could regulate steady state levels of different transcripts to control the timing and expression of key proteins. Models with this type of gene regulation already exist within the scope of reproductive biology: gametogenesis and the post-fertilization maternal-to-zygotic transition. Chromatin condensation occurs throughout both spermatogenesis and oogenesis, with complete transcriptional silencing in mature spermatids and oocytes [53-56]. Consequently, mRNAs are commonly regulated by cytoplasmic polyadenylation to control poly(A) tail length, recruitment of the poly(A) binding protein (PABP), and formation of the translation initiation complex [57-59]. However, additional RNA-BPs provide further regulation. One example is the Deleted-in-Azoospermia protein (DAZ) protein and its autosomal homolog, DAZ-like protein (DAZL). Both DAZ and DAZL recognize target sequences within the 3’ UTR of select mRNAs and can recruit PABP through protein-protein interactions, allowing formation of the initiation complex independently of a poly(A) tail [60-62]. However, the stoichiometry and relative spacing of DAZ/DAZL molecules on a 3’ UTR can recruit repressor proteins (DAZAP, PUM2) whose activity is dependent on phosphorylation status [63-65]. Genes such as the meiosis-associated *sycp3* are regulated by all of the aforementioned regulatory layers, but the net result is a process of ordered and synchronized gene expression independent of changes in transcription [61, 66, 67]. In this context, NP1 and CIRBP may be playing critical roles in chromatin remodeling and controlling mRNA stability and/or translation, respectively.

In mammalian systems, CIRBP plays many diverse, yet highly integrated roles. It is classically recognized for stress granule formation in response to cellular stressors, including heat shock, UV irradiation, hypoxia, oxidative stress osmotic shock, or arsenic exposure [47, 68]. CIRBP is normally stored in the nucleus, and only translocates to the cytosol upon stress induction where it binds target mRNAs and associates into stress granules [47]. As “cold inducible” RNA binding protein, CIRBP received its name because of its role in testes, which operate ~3°C below normal human body temperature [69]. It was later identified as a major cold shock protein in *Arabidopsis*, and the *Arabidopsis* homolog could functionally replace non-homologous cold shock proteins in *E. coli* [70]. The temperature-dependent expression is a product of alternative promoter usage, producing a transcript with an elongated 5’ UTR that contains an internal ribosome entry site [71]. CIRBP has recently been identified as a candidate gene for temperature-dependent sex determination in the common snapping turtle [72]. Echoing the linkage between the mental gland and cancerous tumors, CIRBP is also part of the human telomerase complex and plays a crucial role in telomere maintenance [73]. Because of the seasonal nature of the mental gland and its presence in a cold-blooded animal, temperature-dependent expression in *P. shermani* seemed like a highly tractable hypothesis. However, 5’ RACE experiments revealed that the CIRBP 5’ UTR length was similar between June and August, and was closer in size to the short mammalian 5’ UTR without the internal ribosome entry site (data not shown). Also, CIRBP IHC revealed exclusively cytoplasmic localization at both time points with relatively uniform labelling, not indicative of stress granules. In one study [48], the low complexity domains of multiple RNA-BPs were found to form hydrogels composed of β-sheet cross strands similar to amyloid fibers. Repeats of the tripeptide Gly/Ser-Tyr-Gly/Ser were determined to be essential for formation, and likely involved π-π overlap between tyrosine residues. Using mutagenesis to compare the effect of 0, 12, 18, 22, or 27 tripeptide repeats, hydrogel size and stability were positively correlated with the number of repeats; in contrast, *P. shermani* CIRBP only has 7 repeats, such that it may form smaller aggregates that are not visible microscopically. Given that CIRBP-RNA interactions induced a conformational change in CIRBP secondary structure and recruited additional CIRBP molecules through cooperativity, it is plausible that CIRBP may coat the entire PMF 3’ UTR (possibly the whole PMF mRNA), and interfere with ribosome binding and translation. Given the dramatic changes in PMF mRNA abundance over gland development, this binding may also facilitate mRNA degradation. Stress granules are generally thought to protect mRNA molecules from degradation [74, 75], but their close proximity and possible association with processing bodies (P-bodies) has led to the hypothesis that there may be mRNA exchange between these macromolecular complexes, leading to mRNA degradation under the proper cellular conditions [76, 77].

Similar to reports on the human homolog [47], we determined that both the RRM and LCD were required for stable, highly cooperative binding of CIRBP to the PMF 3’ UTR. Structural comparison of different RRM-RNA complexes from other RNA-BPs has revealed that the RRM is a highly plastic structure with respect to nucleotide binding. The αβ sandwich structure contains two conserved regions of 7 and 6 amino acids (termed RNP1 and RNP2) which bind single stranded dinucleotides on either DNA or RNA through a range of interactions, including π-π overlap between aromatic residues and the nitrogen bases, hydrophobic interactions between additional aromatics and the sugar moieties, and/or salt bridges with the phosphate between the two nucleotides. Despite these conserved structural elements, sequence specificity is dictated by less conserved residues forming additional interactions [78]. RNA secondary structure also plays a critical role, with some RNA-BPs requiring dinucleotides to be part of stem loops or other types of internal loops [79-81]. When the dinucleotide frequency within the PMF 3’ UTR was examined relative to predicted values based on nucleotide abundance, five of the sixteen combinations were observed at higher than expected frequencies: UC (+26%), UG (+58%), CA (+7%), CU (+29%), and GA (+8%). When these distributions were compared by χ2 test, UG contributed much more to the test statistic relative to the other dinucleotides with higher than expected frequencies (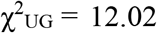, compared to 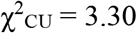). Future experiments examining both CIRBP-RRM specificity as well as PMF 3’ UTR secondary structure will be required to determine if these altered frequencies hold functional significance.

In addition to CIRBP, many other cold shock proteins were identified in the mental gland and had similar expression patterns (Figs 4, S2). All evidence suggests that CIRBP likely plays some critical role in facilitating the glands transition from growth and development into pheromone synthesis by interacting with PMF and likely other mRNAs for an unknown function (likely relating to translational repression and/or mRNA degradation). As already noted, while CIRBP was originally characterized based on its roles in temperature-dependent expression, more recent studies have demonstrated its roles in systemic and general stress response [47, 82]. When we also include factors like NP1, and the transcription factors ATF3 (found in Cluster 1) and NRF2 (possibly present, based on identification of its inhibitor Keap1), the mental gland seems to express a preponderance of stress response genes [83, 84]. It is worth highlighting that this yet another feature the mental gland shares with many cancerous tumors [85, 86]. In addition to facilitating proliferation and development, the role of CIRBP as a regulator of pheromone synthesis is also interesting within the scope of gene co-option. A common theme among plethodontid pheromone genes– as well snake toxins and the products of other exocrine tissues – is gene duplication followed by rapid evolution which drives neofunctionalization [32, 87, 88]. Through the same basic processes, it is plausible that novel regulators could be recruited and co-opted to provide tight control over gene expression of exocrine products in these systems. With many of these gene products having evolutionary histories of positive selection, it would likely be of value to explore both the products and their regulators in a co-evolutionary framework. At least in the case of PMF, CIRBP may have been the causative agent driving purifying selection on the UTRs while sexual selection from the female receptors promoted positive selection on the coding regions. The putative role of CIRBP directing the selective forces on the noncoding regions of PMF might be relevant to future studies of the regulatory elements of other rapidly evolving genes.

## Conclusions

For more than 60 million years, plethodontid salamanders have utilized a rapidly evolving system of non-volatile proteinaceous courtship pheromones that regulate female mating behavior and facilitate reproduction. In the red-legged salamander (*P. shermani*), these pheromones are synthesized in a submandibular mental gland that annually hypertrophies into a large, pad-like structure whose transcriptional and translational machinery are almost exclusively programmed for pheromone synthesis. We now report that this highly effective system results from a tightly coordinated gene expression cascade, allowing for annual organogenesis followed by rapid conversion to a highly efficient pheromone factory. Glandular cells initially have an almost cancerous phenotype characterized by rapid proliferation and ECM dissolution, followed by a tremendous increase in pheromone mRNA levels. A key regulator in this process is Cold Inducible RNA Binding Protein: a stress-responsive RNA binding protein used by both animals and plants to store select mRNAs in stress granules and promote cell survival. For at least one pheromone (Plethodontid Modulating Factor), CIRBP selectively binds the 3’ UTR and recruits additional molecules through cooperativity, with protein-protein and protein-RNA interactions likely stabilized through induced intermolecular β-sheets. This interaction may inhibit translation of PMF mRNA and/or promote its degradation through association with P-bodies. The net result is suppression of pheromone translation until the gland is sufficiently large to support the storage of 10s to 100s of micrograms of pheromone, which can be used to increase receptivity in almost any female in the breeding population. CIRBP may be one player that has exerted purifying selection on the PMF UTRs, creating a system of disjunctive evolution with highly conserved noncoding segments and rapidly evolving coding regions. The mechanisms behind this exciting dichotomy may serve as a model for future studies of gene regulation on rapidly evolving proteins.

## Materials and Methods

### Animal collection, gland removal, and pheromone extraction

*P. shermani* males were collected during their breeding season from a single site in Macon Co., North Carolina, USA (35°10’48” N, 83°33’38” W). Males were anesthetized in a mixture of 7% (v/v) diethyl ether in water. For analysis of total RNA, pheromone extract, or cellular proteins, mental glands were surgically removed using iridectomy scissors. For RNA analysis, glands were incubated overnight in RNAlater (Ambion, Austin, TX) at 4°C before long term storage at -20°C. Pheromone was extracted from mental glands based on the protocol in Chouinard et al. [33]: briefly, mental glands were individually incubated in 0.2 mL acetylcholine chloride (0.8mM in Amphibian Ringer’s solution) for 30 min, centrifuged at 14,000 *x g* for 10 min, the supernatant collected, and centrifugation repeated before storage at -80°C. Following extraction, mental glands were stored in RNAlater to allow preservation of RNA and cellular proteins. Pilot experiments confirmed that 30 min in the acetylcholine solution was sufficient to extract >90% of the total pheromone with no detectable RNA degradation (Wilburn and Feldhoff, unpublished data). Salamanders were collected under permits issued by the North Carolina Wildlife Resource Commission (to R.C. Feldhoff and D.B. Wilburn), and all animal protocols were approved by University of Louisville IACUC (#12041 to Pamela W. Feldhoff) and Highlands Biological Station IACUC (to D.B. Wilburn).

### cDNA preparation and transcriptome sequencing

Mental glands were collected from male *P. shermani* at six time points approximately every 3 weeks during 2010 (5/29, 6/19, 7/10, 8/1, 8/21, and 9/11). This range preceded and spanned the principal August mating season. Five glands were collected at each time point and immediately stored in RNAlater. Total RNA was extracted from individual glands using the RNeasy kit (Qiagen, Valencia, CA) following the manufacturer’s instructions. RNA concentrations were estimated by 260 nm absorbance. For each time point, standardized amounts of RNA were pooled from all five glands, and double-stranded cDNA prepared by oligo-dT priming using the SMARTer cDNA synthesis kit (Clontech, Palo Alto, CA). cDNA was supplied to Otogenetics Corporation (Norcross, GA) for library preparation and sequenced using the Illumina HiSeq 2000 platform (>20 million reads per time point, 100-bp paired end reads). Data was deposited in the NCBI Sequence Read Archive (BioProject ID SUB3398845).

### Transcriptome bioinformatics analysis

Illumina reads from all six time points were pooled and assembled into a single transcriptome using Trinity (r2012-10-05) [89]. Initial assemblies with default settings resulted in over-compaction of deBruijn graphs for PMF, limiting both isoform detection and full-length mRNA re-construction. Butterfly parameters were then optimized, and the final assembly included the additional settings *–min_kmer_cov 2 –bfly_opts “-path_reinforcement_distance= 25 -min_per_id_same_path= 98”*. Reads from each time point were re-aligned to the full transcriptome using RSEM (v1.2.5) [90], with differential expression analysis conducted using EBSeq [91]. Expression differences were compared between adjacent time points (5/29 to 6/19, 6/19 to 7/10, etc.). Based on visual observations and analysis of pheromone extract from additional glands collected at each time point, the separate phases of gland development were best characterized by the 6/19 time point (growth/development) and the 8/1 time point (pheromone production); therefore, an additional comparison with EBSeq was performed between the 6/19 and 8/1 time points. For putative gene annotation, the Trinotate package (r2013-02-25) was used to determine (1) putative open reading frames (using TransDecoder, [89]), (2) protein BLAST (blastp) against both the SwissProt and TrEMBL databases [92-95], (3) orthologous group identification with eggNOG [96], (4) gene ontogeny assignment [97], (5) signal peptide prediction with SignalP [98], (6) protein family assignment with Pfam [99], and (7) transmembrane domain prediction with TmHMM [100]. As most of these databases searches relied on proper open reading frame assignment by TransDecoder, an additional BLAST search was performed with blastx using assembled nucleotide sequences against the full TrEMBL database [93, 95]. There were multiple cases of the TransDecoder proteins having no blastp hits in SwissProt or TrEMBL, yet the nucleotide sequence produced strong blastx hits (E-value < 0.001), suggesting that TransDecoder identified the wrong open reading frame. In these cases, an alternative open reading frame was selected based on the longest amino acid sequence that contained the aligned region of the blastx hit.

### qRT-PCR analysis of differentially expressed genes

For select genes (see Table S4 for primers), qRT-PCR analysis was performed on RNA isolated from each mental gland used to construct the transcriptome (six time points each with 5 glands, 30 glands total). Total RNA was diluted to ~5-20 ng/μL, and accurate concentrations were determined using Quant-iT RiboGreen RNA assay kit (Invitrogen, Carlsbad, CA). qRT-PCR reactions were performed in triplicate using 20 μL reactions with the Power SYBR Green RNA-to-CT 1-Step kit (Ambion) containing 1 μL diluted total RNA. Expression levels were calculated by the pcrfit function in the R package *qpcR* using the cm3 mechanistic model [101]. Based on the gross morphological changes of the mental gland, it was expected that few genes would be stably expressed across mental gland development; therefore, RNA input was used to normalize qPCR measurements, with the literature supporting that input is often a more robust reference than housekeeping genes [102, 103].

Expression levels for each gene were normalized based on the time point with the highest expression, and a single linear mixed effect model was fit by maximum likelihood using the lme function in the R package *lmer*. The model included fixed effects for gene and time, with male as a random effect; all variables, including the interaction between gene and time, were significant at p < 0.001.

### Expression of recombinant CIRBP

To conduct *in vitro* protein-RNA binding assays, recombinant CIRBP was prepared using an *E. coli* expression system. Preliminary experiments revealed that CIRBP was insoluble at > 0.2 mg/mL in all tested buffers (data not shown); however, solubility was dramatically improved when recombinant CIRBP was expressed as a fusion protein with enhanced cyan fluorescent protein (rCIRBP/ECFP). Simultaneously, fusion proteins were prepared with only the RRM-containing N-terminus (residues 1-84; rCIRBP-RRM/ECFP) and the glycine-rich C-terminus (residues 85-165; rCIRBP-LCD/ECFP). All constructs included ECFP on the N-terminus, a short hydrophilic linker (SGAAAAGGSDP), and the CIRBP element at the C-terminus. The CIRBP coding regions were amplified from mental gland cDNA using the Accuprime High Fidelity *Taq* polymerase system (Invitrogen) (see Table S4 for primers). ECFP was amplified from a pcDNA3.1-based vector (supplied by Dr. Ronald Gregg, University of Louisville), and fusion genes were prepared by modified assembly PCR [104]. Fusion PCR products were purified using the QIAquick PCR cleanup system (Qiagen), and cloned the pET45b vector (Novagen, San Diego, CA) following restriction digest with *Kpn*I and *Hind*III (New England Biolabs, Ipswisch, MA), gel purification of cleavage products (GFX purification system, GE Healthcare, Piscataway, NJ), ligation using T4 DNA Ligase (New England Biolabs), and transformation into T7 Express lysY/Iq chemically competent *E. coli* (New England Biolabs). Sequence of transformed DNA was validated by colony PCR and Sanger sequencing by the University of Louisville DNACore facility. For protein expression, clones were cultured overnight at 28°C in LB media with 100 μg/mL ampicillin (LB/Amp), diluted in 1 L LB/Amp with a 600 nm optical density (OD600) equal to ~0.05, incubated with shaking until the OD600 equaled ~0.7, IPTG added to a final concentration of 0.1 mM, and incubated overnight. *E. coli* were harvested by centrifugation, resuspended in 50 mM NaCl/0.1% Triton X-100/2 mM EDTA/50 mM Tris, pH 8, and lysed by sonication followed by lysozyme treatment (final concentration 1 mg/mL) for 1 hour. Insoluble material was removed by centrifugation, and proteins concentrated by ammonium sulfate precipitation (final concentration 70%). Ammonium sulfate pellets were resolubilized in Ni-NTA binding buffer (0.5 M NaCl/5 mM Imidazole/20 mM Tris, pH 8), and passed through a 15 mL Ni-NTA column (Thermo-Pierce, Rockford, IL) at 1 mL/min. The column was then washed and eluted with increasing concentrations of imidazole (all in 0.5 M NaCl/20 mM Tris, pH 8): 10 column volumes (CVs) at 5 mM, 2 CVs at 20 mM, 1 CV at 40 mM, 1 CV at 60 mM, and finally 3 CVs at 200 mM (collected in 5 mL fractions). Highly fluorescent fractions were pooled, concentrated, and buffer exchanged to 1X EMSA Buffer (100 mM KCl/2 mM EDTA/0.05% Tween-20/20 mM HEPES, pH 7.4) by ultrafiltration (YM-10 Centiprep, Millipore). All fusion products were standardized to equal molar concentrations (rCIRBP/ECFP: 1.4 mg/mL, rCIRBP-N/ECFP: 1.1 mg/mL, rCIRBP-C/ECFP: 1.0 mg/mL).

### Electrophoretic mobility shift assays

CIRBP-RNA interactions were characterized by electrophoretic mobility shift assays (EMSAs). RNA based on the PMF Class I 3’ UTR was prepared by *in vitro* transcription using the T7 High Yield RNA Synthesis Kit (New England Biolabs) with purified PCR products amplified from *P. shermani* 8/1 cDNA using primers that included engineered T7 promoters (Table S4). Synthesized RNA was subsequently treated with Turbo DNase I (Ambion), and purified using the RNeasy Kit (Qiagen). For fluorescently labeled RNA, *in vitro* transcription reactions were adjusted to include 7.5 mM UTP and 2.5 mM aminoallyl-UTP (Ambion), treated with TURBO DNase I (Ambion), purified using the RNeasy kit (Qiagen), and adjusted to 0.4 mg/mL with 2.5 mg/mL TAMRA-carboxylic acid (Invitrogen) in 0.1 M MES (pH 5). One-fourth volume of EDAC (0.1 mM) was added to the reaction, incubated for 2 hours in the dark, and then labelled RNA was purified using the RNeasy Kit (Qiagen). All EMSAs were performed as 15 μL reactions in 1X EMSA Buffer with 3% glycerol using either 300 ng unlabeled or 45 ng labeled RNA. rCIRBP/ECFP (or the domain truncations), competitors, and other reagents were added at the listed concentrations, and incubate at room temperature for 20 minutes prior to gel loading. For crosslinking assays, formaldehyde was added to the protein-RNA mixture at a final concentration of 1%, incubated for 5 min, and the reaction quenched by addition of glycine to a final concentration of ~330 mM prior to gel loading. RNA was separated using agarose gels of 2% (for PMF 3’ UTR 26-667) or 3% (for ~250 nt products), and electrophoresis performed at 80 V for 3 hours. For unlabeled RNA, gels were stained with Sybr Green II RNA Stain (Invitrogen) for 30 min. Gels were imaged using a Typhoon 9400 fluorescent bed scanner (GE Life Sciences, Piscataway, NJ) with the appropriate laser and filter settings. For fluorescence quenching assay, 0.5 μg of TAMRA-labeled RNA was incubated with increasing concentrations of different rCIRBP constructs in a 200 μL reaction, and fluorescence measured using the Synergy2 plate reader (Biotek, Winooski, VT). Data were fit to the Hill equation (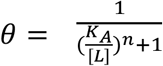) using the nlsLM function in the R package *minpack.lm*.

### Antibody preparation

PRF antisera was prepared by immunizing two rabbits with PRF fractions collected by anion-exchange chromatography (performed by RCF at U. Louisville). The antigen was >90% enriched for PRF, but contained ~5% PTP such that there was significant immunoreactivity for both proteins. CIRBP antisera was prepared by immunizing two rabbits with purified rCIRBP/ECFP (~98%) by Ni-NTA chromatography (performed by the Proteintech Group, Chicago, IL). Highly purified antigens (>99% by reverse phase HPLC) were coupled to agarose matrices for affinity purification of antibodies: briefly, 6% crosslinked agarose beads (CL6B; Sigma-Aldrich, St. Louis, MO) were activated with carbonyldiimidazole (CDI; Sigma-Aldrich) in dry acetone for 20 minutes, quickly rinsed under vacuum with cold distilled water, and then incubated in antigen solutions (~2 mg/mL in 100mM KCl/0.05% Tween-20/100mM NaCO_3_, pH 11) overnight with gentle agitation. Resins were packed into ~1-3mL columns (depending on the starting amount of antigen), remaining active sites blocked with 50 mM Tris, pH 10, and equilibrated in 1X phosphate buffered saline (PBS). Antibodies were purified by incubating antisera with the resin overnight at 4°C, washing with 10 CVs of 1X PBS to collect unbound antibodies, stringently washed with 5 CVs of 0.5 M NaCl/0.05% Tween-20/20 mM Tris, pH 7.5 (TTBS), and eluted with 6 CVs 0.1 M citric acid/0.1% Tween-20, pH 3 collected in 0.2 mL fractions. Elution fractions were neutralized with 1 M Na_2_HPO_4_, and antibody-containing fractions determined by absorbance measurements at 280 nm. Fractions were then pooled, concentrated, and buffer exchanged to 1X PBS by ultrafiltration (YM-30 Centriprep; Millipore, Billerica, MA). Four antigen columns were prepared: PRF, PTP, rPRF/ECFP, and rCIRBP-RRM/ECFP. PRF and PTP antibodies were serially purified from the same antisera by first passing whole antisera through the PRF column for purification of anti-PRF, and the flow through (i.e., unbound antibodies) then passed over the PTP column for purification of anti-PTP. Both solutions of affinity-purified antibodies were then adsorbed against the opposing column to ensure removal of non-specific or low-affinity antibodies. anti-CIRBP-RRM was purified by passing rCIRBP/ECFP antisera over the rPRF/ECFP column to remove anti-ECFP, the flow through next incubated with the rCIRBP-N/ECFP column, anti-CIRBP-RRM purified, and the two-column process repeated to ensure removal of non-specific or low-affinity antibodies. Antibody specificity was validated by western blot.

### Histological analysis

For histological analysis of mental glands, male salamanders were anesthetized in 7% ether, sacrificed by rapid decapitation, the lower jaw removed, immediately placed in 4% paraformaldehyde (in 150 mM NaCl/100 mM Na_2_HPO_4_, pH 7.4), and incubated overnight with gentle agitation. Fixed jaws were decalcified by incubation in 10% EDTA (pH 7.4, DEPC-treated) for 48 hours, cryoprotected in 30% sucrose for 48 hours, embedded in Optimum Cutting Temperature (OCT) media (Sakura-Finetek, Torrance, CA), and stored at - 80°C prior to cryosectioning. Lower jaws were sectioned coronally at a thickness of 16 μm (for immunohistochemistry) or 40 μm (for structural morphology), and thaw mounted onto superfrost plus slides pre-coated with polylysine. Slides were stored at -20°C until analyzed. Hematoxylin/eosin staining was performed using standard protocols [105]. For immunohistochemical labeling, slides were first equilibrated to room temperature for 30 min and washed five times with 1X PBS for 5 min each. Antigen retrieval was conducted by incubating the slides in 10mM citric acid (pH 6)/0.05% Tween-20 at 70°C for 30 min, cooled to room temperature for 20 min, and washed twice with PBS with 0.05% Tween-20 (PBST) for 5 min each. Blocking was performed for 30 min with 1X PBS/0.1% BSA/0.5% Tween- 20, and incubated overnight with primary antibody (anti-PRF at 7.5 μg/mL or anti-CIRBP-RRM at 3.5 μg/mL) in PBST with 0.1% BSA. Slides were then washed five times with PBST for 5 min each, incubated with secondary antibody (Alexa Fluor 633 goat anti-rabbit IgG (Invitrogen), 1 μg/mL in PBST) for 30 min, washed five times with PBST for 5 min each, and finally counterstained with Alexa Fluor 488-wheat germ agglutinin (Invitrogen; 10 μg/mL in PBST) and DAPI (Invitrogen; 3.6 μM). For gland morphology, thick sections (40 μm) were stained with Alexa Fluor 488-wheat germ agglutinin, Alexa Fluor 555-Phalloidin (Invitrogen; 0.005 U/μL), and DAPI. Slides were cover slipped with Prolong Gold Antifade reagent (Invitrogen), cured overnight in the dark, and stored at 4°C. Imaging was accomplished using an Olympus Fluoview FV1000.

### Circular Dichroism

To measure secondary structure changes in rCIRBP/ECFP with different concentrations of RNA, far-UV CD spectra (185-255 nm) were acquired by averaging 10 scans across a 0.1-cm path at 0.2 nm intervals using a Jasco J-810 Spectropolarimeter. Stock rCIRBP/ECFP (1.4 mg/mL in 1X EMSA) buffer was diluted 10-fold in water (to reduce chloride background). Following initial measurements with no RNA, PMF 3’ UTR 26-667 (at 70 ng/μL in 0.1X EMSA Buffer) was titrated into the solution, incubated for 5 min, and CD spectra recorded. Spectra were adjusted for slight changes in volume/concentration following serial dilution throughout the experiment. At the concentrations used, PMF 3’ UTR 26-667 produced no significant CD signal over buffer.

### Quantification and modelling of CIRBP expression

For two separate time points in 2013 (6/13 and 8/3), 15 male *P. shermani* were collected, mental glands removed, pheromone extracted, and mental glands stored in RNAlater (Ambion) as previously described. One gland for 8/3 was inadvertently destroyed, such that n = 29. Pheromone concentrations were accurately determined by BCA Protein Assay (Thermo-Pierce), and normal proportions of pheromone components (PMF:PRF:PTP ~ 5:3:1) were validated by RP-HPLC. Stored mental glands were later dissected into approximately equal halves to extract both RNA (RNeasy Kit, Qiagen) and cellular protein (homogenization in RIPA Buffer, supplemented with 0.1 mM iodoacetamide). Pilot studies confirmed that the two halves were equivalent for levels of target proteins and mRNA. CIRBP protein levels were measured by western blot. Preliminary experiments suggested that glands from 6/13 had ~6X higher levels of CIRBP, such that to maintain similar blot intensities for quantification, 5 μg was loaded per lane for 6/13 glands and 30 μg per lane for 8/3 glands. Briefly, proteins were separated using 15% Tris-Tricine gels with 4% stacking gels [106] at 50 V for 15 minutes followed by 100 V for 90 min, transferred to PVDF membranes by electroblotting in 15% methanol/0.025% SDS/10 mM CAPS, pH 11 at 55 V for 75 min. The membrane was blocked with 0.5 M NaCl/0.5% Tween-20/20 mM Tris, PH 7.5 for 1 hour, incubated with anti-CIRBP-RRM (1.4 μg/mL) for 1 hour, then incubated with alkaline phosphatase-conjugated goat anti-rabbit IgG (1 μg/mL; Sigma-Aldrich), and developed using BCIP/NBT. All membrane washes and antibody dilutions were performed using TTBS (0.5 M NaCl/0.05% Tween-20/20 mM Tris, pH 7.5). All blots were processed and developed simultaneously to minimize run-to-run variation, and a reference of 10 ng rCIRBP/ECFP was included on each membrane for normalization. Densitometry analysis was performed using ImageJ. Extracted RNA was used for qRT-PCR analysis for PMF, CIRBP, and Cathepsin S based on the previous methods (see Table S4 for primers). Correlation between variables was determined by multivariate ANOVA (MANOVA).

### Immunopulldown and mass spectrometry of CIRBP

To validate the identity of different bands visualized during western blot, immuno-pulldown experiments were performed. Anti-CIRBP-RRM was used for CIRBP pulldown, and as a negative control, an IgG-enriched fraction from pre-immunization serum of CIRBP immunized rabbits was prepared by ammonium sulfate precipitation and DE52 chromatography. Using Protein G coupled Dynabeads (Invitrogen), 10 μg of antibody was adsorbed to the beads, stringently washed with PBST, incubated with pooled RIPA extract from 6/13 (~180 μg) or 8/3 (~840 μg) for 30 min, washed with PBST, the beads transferred to a clean 1.7 mL tube, and incubated with 1X gel loading buffer (1% SDS/12% glycerol/0.005% bromophenol blue/50 mM Tris, pH 6.8) at 65°C for 30 min. The complete samples were loaded into a 15% Tris-Tricine gel with 4% stacking gel, electrophoresed at 50V for 15 min followed by 100 V for 90 min, and stained using SYPRO Ruby fluorescent gel stain (Invitrogen). The gel was imaged using the Typhoon 9400 fluorescent bed scanner (GE Life Sciences). All bands were then individually excised, proteins reduced and alkylated with dithiothreitol/iodoacetamide, treated overnight with trypsin, peptides isolated by acetonitrile extraction, and supplied to the University of Louisville Biomolecular Mass Spectrometry Core Laboratory for analysis by LC/MS-MS.

## Authors′ contributions

The study was conceived and designed by DBW and RCF. All experiments and data analysis were performed by DBW. The manuscript was drafted by DBW and edited by RCF.

## Acknowledgements

We thank Pamela W. Feldhoff, Kari A. Doty, Kathleen Bowen, Lynne D. Houck, Stevan Arnold, Sarah Eddy, Adam Chouinard, and Sean Darrow for assistance in the field, and the Highlands Biological Station for providing laboratory space and housing. We also thank the University of Louisville Biomolecular Mass Spectrometry Core Laboratory for their continued support. Funding was provided in part by National Science Foundation (Collaborative) grants IOS-1146899 (RCF), a NSF Graduate Research Fellowship to DBW, and two HBS Grant-in-Aid awards to DBW.

